# Graphene oxide chemically decorated with hybrid Ag-Ru/chitosan nanoparticles: fabrication and properties

**DOI:** 10.1101/022400

**Authors:** Murugan Veerapandian, Suresh Neethirajan

**Affiliations:** BioNano Laboratory, School of Engineering, University of Guelph Guelph, ON N1G 2W1, Canada

**Keywords:** Hybrid graphene oxide, Ag-Ru complex, Chitosan, Functionalization, Biosensor platform

## Abstract

Hybridization of distinct materials into a single nanoplatform is relevant to advance material’s properties for functional application such as biosensor platform. We report the synthesis and characterization of nanosheets of graphene oxide decorated with hybrid nanoparticles of silver-ruthenium bipyridine complex (Ag@[Ru(bpy)_3_]^2+^) core and chitosan shell. Hybrid nanoparticles were first obtained through a sequential wet-chemical approach using *in situ* reduction, electrostatic and coordination reaction. Oxygenated functional groups of graphene oxide and abundant amine groups of chitosan layer on the surface of hybrid nanoparticles allowed the functionalization reaction. Changes in intrinsic optical, chemical and structural properties of graphene oxide due to hybrid nanoparticles were studied in depth using spectroscopic techniques and an electron microscope. Electrodes modified with nanosheets of graphene oxide-hybrid nanoparticles retain the biocompatibility and displayed an amplified redox property suitable for a broad range of sensing studies.

## 1. Introduction

Research on the development of hybrid nanomaterials with multiple structures and chemical composition has attracted the attention of many researchers to advance the functional properties [1]. In recent years, two-dimensional graphene oxide (GO) and reduced GO (rGO) have been used in a variety of applications due to their cost-effective fabrication, ultra-thin layers, large surface area and tunable oxygen functional groups [2-3]. Surface treatment and functionalization of active components on GO nanosheets influence the inherent sp^2^/sp^3^ carbon domains, which mediate the change of crystallite size, lattice orientation and associated physico-chemical properties [4].The different strategies employed to tune the physico-chemical and biomedical functionality of GO are photoirradiation, elemental doping and chemical anchoring of inorganic/organic materials [3-6].

Significant effort has been put into the development of advanced hierarchical nanostructures based on hybrid GO materials. In particular, ternary/quaternary nanocomposite comprised of graphene-derivatives, metal, metal-oxide and polymer has recently been shown to have improved physico-chemical properties optimal for device construction (e.g., electrode materials for biosensor platform and energy conversion). Typical enhanced electrochemical properties of metalloid polymer hybrid (Ag@SiO_2_-PEG)-GO [7], graphene/WO_3_/Au [8] and polyaniline-Fe_2_O_3_-rGO [9] are well-suited to biosensor studies than the individual pristine derivatives. Molecularly imprinted polymers based on CdTe/Cds and magnetic GO showed selective recognition toward environmental pollutants [10]. Pt-graphene-TiO_2_ [11] and reduced GO-bismuth ferrite (Bi_2_Fe_4_O_9_) [12] have been reported to have better photocatalytic properties. Further, studies show that the hierarchical structures of SnS_2_-rGO-TiO_2_/TiO_2_ layered films [13] and rGO/Fe_3_O_4_@SiO_2_@polyaniline [14] significantly improve photoelectric and electrochemical properties, respectively. The accumulation of evidence indicates that the fabrication of hybrid GO material has great potential for opto/electrochemical device development.

Nevertheless, achieving a durable structure of hybrid GO with inbuilt multi-functionality is complicated. Physically-linked hybrid nanostructures are prone to leaching and decreased synergistic functionality. Compared to physical adsorption of nanostructures on GO surface, chemically-bonded active materials can retain better stability and are expected to have durable electrochemical properties. However, only few studies have demonstrated the chemical functionality between active materials and GO surface. Incorporation of a durable single hybrid nanostructure on GO surface with optical, electrochemical and biocompatible capabilities would be highly useful for various biosensing applications. Previous work has shown that single hybrid core-shell nanoparticles made of a metal-dye complex (AgNPs@[Ru(bpy)_3_]^2+^) core and biopolymer (chitosan) shell can influence optical, electrochemical and biocompatibility due to the electrical conductivity of Ag, metal-to-ligand charge-transfer of [Ru(bpy)_3_]^2+^ and abundant amino groups of chitosan [15]. The three-in-one hybrid [15] nanosystem (average particle size 54 nm) on the surface of GO would be an ideal candidate for modification; it is multifunctional due to its opto-electronic and biocompatible nature. GO chemically decorated with hybrid nanoparticles (HNPs) is expected to have better optical, redox activity and biocompatible functional groups suitable for various sensor studies. For example, introduction of Ag metal on GO improves the electron transfer process and increases immunosensing ability [16]. The presence of metallic and hydrated Ru on the surface of GO electrodes enhanced electrochemical performance [17,18]. As a bio-derived linear polysaccharide with biocompatibility, biodegradability and film-forming ability, chitosan is explored as an interface layer in the fabrication of chemically-modified electrodes for biosensing [19,20].

Chemical functionalization of distinct materials on the surface of GO is highly dependent on the reactivity of oxygen functional groups which exist on the edges and basal planes of GO. Here, for the first time, the three-in-one HNPs of Ag@[Ru(bpy)_3_]^2+^/chitosan are used to chemically decorate the GO nanosheets. Abundant amino groups of chitosan-coated on the surface of Ag@[Ru(bpy)_3_]^2+^ provided a significant modification on the oxygenated edges/basal planes of GO. The influence of optical absorbance, photoluminescence, zeta potential and structural integrities including morphology, chemical structure and Raman shift of pristine GO and HNPs-GO materials were extensively studied to understand the properties. To evaluate their electrochemical properties, cyclic voltammetric measurements were performed on the customized electrode modified with HNPs-GO. Inherent synergistic physico-chemical properties with biocompatible functional groups derived from the HNPs-functionalized GO may help to construct advanced electrochemical active sensor platforms.

## 2. Experimental Section

### 2.1 Chemicals

Silver nitrate (AgNO_3_), 3-mercaptopropionic acid (3-MPA), sodium borohydride (NaBH_4_), tris(2,2’-bipyridyl)dichloro ruthenium(II) hexahydrate, chitosan (low molecular weight: 50 000-190 000 g mol^−1^; degree of deacetylation: 75-85%), graphite powder (<20 μm, synthetic) and phosphate buffered saline (PBS) were purchased from Sigma-Aldrich. Other chemicals were of analytical grade and used as received without any further purification. Milli-Q water (18.2 MΩ) was used for all experiments.

### 2.2 Synthesis of hybrid (Ag@[Ru(bpy)_3_]^2+^/chitosan) NPs

Hybrid NPs of Ag@[Ru(bpy)_3_]^2+^/chitosan were prepared according to the reported procedure [15]. At first, 5 mL of AgNO_3_ (0.1 M), 25 mL of 14 N aq. NH_4_OH and 5 mL of 3-MPA (50 mM) were dissolved in 15 mL of deionized (DI) water (solution A). Separately, 5 mL of NaBH_4_ (0.02 M) and 2 mL of 14 N aq. NH_4_OH were dissolved in 15 mL of DI water (solution B). At room temperature, solutions of vial A and B were slowly injected dropwise into the 300 mL of DI water over 30 min with a magnetic stirring of 600 rpm. After 30 min of reaction time, colloidal solution containing AgNPs-modified 3-MPA was separated by centrifugation (13,000 rpm for 1 hr). The particles were then washed twice with DI water and dispersed in DI water for further reaction.

[Ru(bpy)_3_]^2+^ coating on AgNPs was achieved by mixing the above AgNPs-modified 3-MPA (5 mL, 1 mg/mL) and ethanolic solution of Ru(bpy)_3_Cl_2_ (5mL, 0.8 mg/mL). The reaction mixture was left overnight under mild stirring, protected from light. Resulted particles were centrifuged (13,000 rpm for 1 hr) and washed twice with ethanol and DI water to remove unreacted [Ru(bpy)_3_]^2+^. Prepared Ag@[Ru(bpy)_3_]^2+^ were then surface modified with chitosan by coordination chemical reaction using Ag@[Ru(bpy)_3_]^2+^ (5 mL, 1 mg/mL) and chitosan (5 mL, 0.01 wt%) under magnetic stirring of 600 rpm for 3 hrs at room temperature. The final hybrid (Ag@[Ru(bpy)_3_]^2+^/chitosan) NPs were isolated by centrifugation (13,000 rpm for 1 hr), washed and re-dispersed in DI water for further experimentation.

### 2.3 Functionalization of HNPs on GO nanosheets

Colloidal dispersions of GO nanosheets used in the current experiment were synthesized according to the modified Hummers’ method [21]. Functionalization of HNPs onto the surface of GO sheets was achieved through a one-step process. An aqueous dispersion of GO (25 mL, 0.5 mg/mL) and HNPs (25 mL, 2 mg/mL) was added to a reaction flask and kept under magnetic stirring (600 rpm) at room temperature for 12 hrs. After the reaction time, the HNPs-functionalized GO sheets were separated by centrifugation (13,000 rpm, 1 hr), washed thrice with DI water and utilized for characterization.

### 2.4 Construction of HNPs-GO sheets modified electrode

An integrated gold printed circuit board (Au-PCB) chip served as the electrode system. The central circle-shaped Au substrate with an area of 2 mm in diameter was used for the modification of HNPs-GO sheets. The two crescent-shaped Au substrates with a length of 4.3 mm and a breadth of 0.8 mm were used as counter and reference electrodes, respectively. After performing sequential washing with acetone, ethanol and DI water, the Au-PCB chip was exposed to plasma treatment. Then, typically 4 μL of the aqueous dispersion of HNPs-GO (1 mg/mL) was drop casted on the working substrate. To make uniform surface modification of HNPs-GO sheets on the electrode surface, typically three layers of casting were performed at regular intervals with an evaporation period of 1 hr at ambient temperature.

### 2.5 Instrumentation

Cary 5000 UV-Vis-NIR spectrophotometer (Agilent Technologies) was used to analyze the UV-vis absorbance spectra. Morphological characterizations were observed via a transmission electron microscope (TEM) (Philips Tecnai 12) with an acceleration voltage of 120 kV. Samples used for imaging were prepared by casting 4 μL of (0.25 mg/mL) HNPs, GO or HNPs-GO suspension onto a carbon-coated nickel grid. Zeta potential was studied from Zetasizer Nano ZS (Malvern Instruments) equipped with a 4 mW, 633 nm He-Ne laser using appropriate cells. Measurements were conducted in backscattering (173°) mode and detected with an Avalanche photodiode. For accurate determination of zeta potential, thirteen runs were averaged for each liquid sample. A Varian Cary Eclipse Fluorescence spectrophotometer was used to examine the photo-luminescence properties of HNPs, GO and HNPs-GO. The chemical structure and functional group modifications on pristine and hybrid materials were identified by Fourier transform infrared (FTIR) spectra studied on a Nicolet 6700 FTIR spectrometer (in the ATR mode, diamond crystal). ^1^H-NMR spectra in deuterated dimethyl sulfoxide-d_6_ (DMSO d_6_) were measured on a Bruker AV 400 spectrometer operating at 400 MHz (number of scan: 256). Raman spectral analysis was performed in RENISHAW inVia Raman microscope equipped with CCD camera and a Leica microscope. Aqueous dispersion of sample (~1 mg/mL) was drop casted on a cleaned silica wafer and utilized for measurements. An excitation wavelength of 514 nm and laser power of 10% was used. A short working distance 50× objective lens was used to focus the laser spot on the sample surface. Measurements were taken in 30s of exposure time at varying numbers of accumulations. Electrochemical properties of the pristine GO and HNPs-GO materials were studied from the cyclic voltammetric technique using SP-150 potentiostat, Bio-Logic instruments. All the cyclic voltammograms (CVs) were recorded in the 10 mM PBS solution (pH 7.4) as supporting electrolyte, in the potential region between 0.25 to +0.8 V. A reproducible voltammogram can be obtained under steady-state conditions after about five cycles.

## 3. Results and Discussion

### 3.1 Synthesis of Ag@[Ru(bpy)_3_]^2+^/chitosan NPs and functionalization on GO

Fig. 1 illustrates the step-wise synthesis route for obtaining HNPs. HNPs made of a metal-dye complex (Ag@[Ru(bpy)_3_]^2+^) core and a chitosan shell are firmly bonded to each other by electrostatic and coordination interaction, respectively [15]. The thin layer of chitosan on the surface of HNPs with abundant amine groups are reactive to the oxygenated functional groups of GO. The presence of carboxyl and epoxyl groups at the edges and basal planes of GO provided multiple binding sites for chemical functionalization of HNPs. The two important surface chemical reactions involved in this functionalization were formation of amidation at the carboxyl groups and nucleophilic attack at the α-carbon by the HNPs. The structural and chemical changes which resulted from the functionalization process were characterized by FT-IR and ^1^H-NMR spectroscopy as described later.

**Fig. 1.**
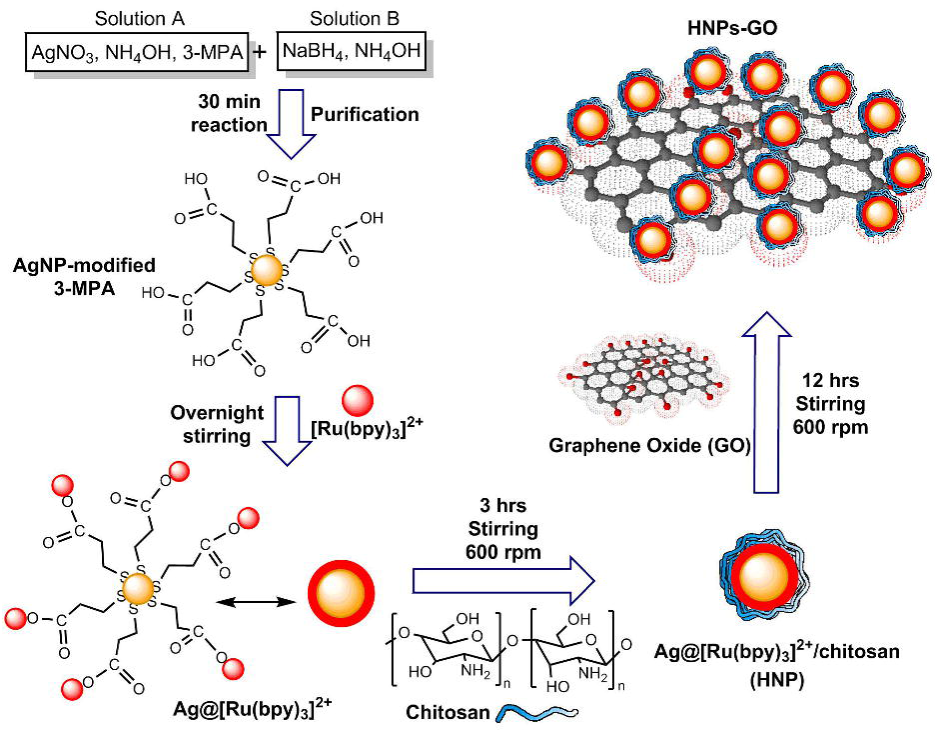
Synthesis of HNP and functionalization on GO nanosheet.

### 3.2 UV-vis absorbance and photoluminescence

In general, the optical absorption and emission band of metal or metal hybrid nanostructures depend on the size, shape, nature of the surface functional layer and solvent environment [22]. Here, an UV-vis absorbance spectroscopy was utilized to measure the optical information of the prepared materials. Fig. 2A represents the spectra observed from aqueous AgNPs and AgNPs-modified with 3-MPA. The peak at 402 nm denotes the existence of characteristic surface plasmon resonance (SPR) of AgNPs. An SPR is the collective oscillations of the conductive electrons that exist on the surface of metal NP. Depending on the excitation of the localized surface plasmon, caused by strong light scattering at a specific wavelength, strong SPR bands are produced [23]. At 423 nm, a significant red shift was mediated by surface modification of AgNPs with 3-MPA.UV-vis absorption spectrum of [Ru(bpy)_3_]^2+^ shows the three specific peaks at 242, 290 and 450 nm (Fig. 2B) ascribed to intra-ligand transition π→π*, bpy π→π_1_* transition and metal-to-ligand charge-transfer (MLCT) band, respectively [24]. A shoulder peak at 420 nm is also attributed to MLCT (t_2g_ (Ru)→π* (bpy) transitions). Similarly, AgNPs modified with [Ru(bpy)_3_]^2+^ also exhibit three absorbance peaks with a moderate hump located at 423 nm due to the overlap of SPR from AgNPs [25].

**Fig. 2.**
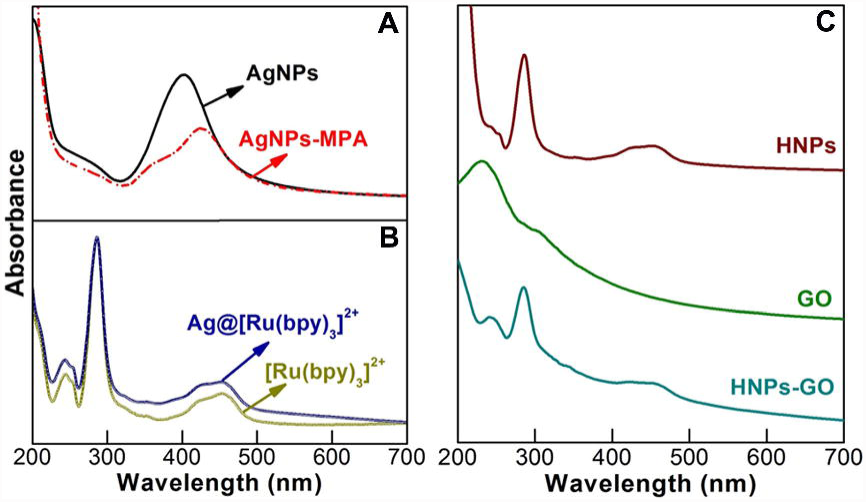
UV-vis absorbance spectra of aqueous dispersion of different materials.

As shown in Fig. 2C, HNPs exhibited significant changes in the peak shape at 242, 290 and 450 nm, indicating that the chitosan modification altered the optical absorbance of Ag@[Ru(bpy)_3_]^2+^ [15]. Aqueous dispersion of GO nanosheets exhibited a wavelength of maximum absorbance at 230 nm attributed to the π→π* electron transition of the polyaromatic C-C bonds of GO layers [7]. The UV-vis absorbance spectrum of HNPs functionalized GO exhibits peaks centered at 240, 284 and 450 nm. Compared to pristine HNPs, the π→π* electron transition signals of HNPs-GO are well resolved, probably due to the associated signals of metal-dye complex and C-C bonds of GO. There is no significant change observed from the bpy π→π_1_* transition peak position, however the peak centered at 450 nm is much more broad than that of pristine HNPs.

Excitation of specific wavelengths of light on the aqueous dispersion of optically active nanomaterials could provide additional information such as photoluminescence (PL). It is known that chemical oxidation of graphite results in the formation of mixed sp^2^/sp^3^ domains in GO lattice, which creates a disruption of the π-network and generates an emission band [26]. The PL spectrum of GO and HNPs-GO was recorded using an excitation wavelength of 325 nm and is shown in Fig. 3. GO shows a sharp emission peak in the near UV region at around 365 nm due to the amorphous sp^3^ matrix that surrounds the various graphitic sp^2^domains, which act as a high tunnel barrier resulting in the generation of a band gap in GO. This is in agreement with previous reports on PL of GO [26,27]. Upon modification with HNPs, the near band emission is quenched with slight broadening of the peak centered at 362 nm. This observation is probably due to the formation of new metallic hybrid clusters on the GO lattice. The observation of PL from HNPs-GO implies the existence of a band gap in the electronic structure of the material. Recent studies identified that the surface modification of GO could create a large band gap and decent carrier mobility suitable for advanced PL [28] and electrochemical biosensor [29]. As-prepared chemically decorated GO containing ternary composite of metal-dye complex and biopolymer retained the inherent PL property and electronic structure, and is expected to be a feasible option for dual (optical/electrochemical) sensors.

**Fig. 3.**
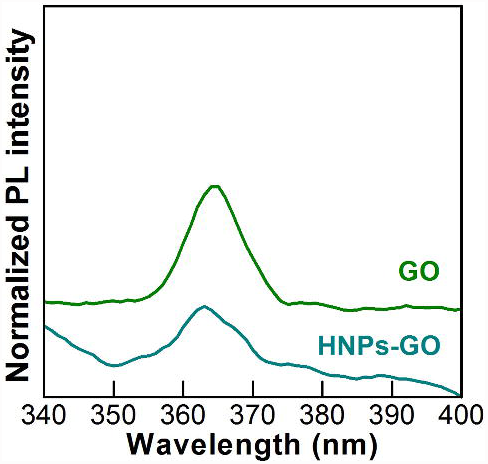
PL spectra of aqueous GO and HNPs-GO nanostructures.

### 3.3 Morphology and surface zeta potential characterization

Morphology of HNPs, GO and HNPs-GO nanostructures were visualized from TEM and are shown in Fig. 4. HNPs with an overall spherical shape and coating of chitosan layer are clearly visible in Fig. 4(A-B). Observed trace of particle’s aggregation is possibly due to the drying process done before imaging. Average particle size distribution of HNPs was determined using the Malvern-dynamic light scattering-Zetasizer Nano ZS instrument and found to be 54 nm (data not shown). Surface topography of GO (Fig. C-D) displays the corrugated thin sheet-like membranous layer. The typical thin grooves or wrinkles on the sheets are characteristic of GO nanostructures. Due to its two-dimensional thin layered feature with reactive oxygenated groups, GO allows multiple chemical bonding with amine-functionalized HNPs. HNPs were well decorated on the surface of GO nanosheets (Fig. 4 E-F).

**Fig. 4.**
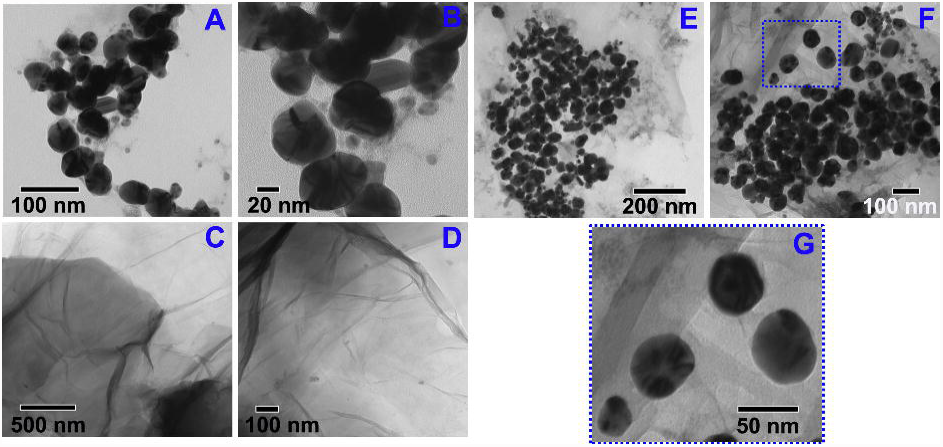
TEM images of (A and B) HNPs, (C and D) GO and (E-G) HNPs-GO nanostructures.

Zeta potential is a vital physical property used to study the stability of colloidal dispersions and surface charge associated with the double layer around the colloidal particle. The zeta potential varies depending on which chemical groups exist on the surface of colloidal particles (supporting Fig. S1). Due to the ionization of the multiple surface oxygenated functional groups, pristine GO showed the negative zeta potential of □39 mV. HNPs containing chitosan shell with abundant amine groups displayed the positive zeta potential of +46.1 mV. Upon surface functionalization, the zeta potential of HNPs-GO was +26.6 mV, indicating that the chemically bonded HNPs modified the inherent surface zeta potential of the GO nanosheets. These results supplement the morphological images. Such modified hierarchical GO sheets with single hybrid of metal-dye complex and biopolymer provide a unique set of physico-chemical properties that are promising for multi-functional material.

### 3.4 FTIR and ^1^H-NMR spectroscopy

In order to evaluate the chemical structure and functional group modifications on HNPs or HNPs-GO, a comparative FTIR spectral analysis was performed on pristine chitosan and GO samples. Fig. 5(A) shows the FTIR spectrum of chitosan: the C-H out of plane bend at 887 cm^−1^ and C-O stretch at 1026 and 1065 cm^−1^. C-O-C stretch and C-H bend are located at 1154 and 1377 cm^−1^. Vibrations at 1590 and 2870 cm^−1^ are attributed to N-H bend and C-H stretch. The broad peak centered at 3317 cm^−1^ is associated with the N-H stretch and hydrogen-bonded OH groups [30,31]. HNPs containing chitosan modified Ag@[Ru(bpy)_3_]^2+^ (Fig. 5B) exhibit significant alterations in their group frequencies. For instance, the C-O stretch shows distinct changes at 1040 cm^−1^ when compared with pristine chitosan. The primary N-H bend at 1590 cm^−1^ is shifted to 1644 cm^−1^, denoting the formation of a secondary amine. A short but sharp peak at 2970 cm^−1^ is ascribed to the asymmetric stretching of C-H [30]. Further, the N-H stretch and H-bonded OH stretch were much more intense than that obtained for chitosan. Observed modifications in the group frequencies (C-O stretch and N-H bend) of chitosan support their chemical bonding with Ag@[Ru(bpy)_3_]^2+^ [15].

**Fig. 5.**
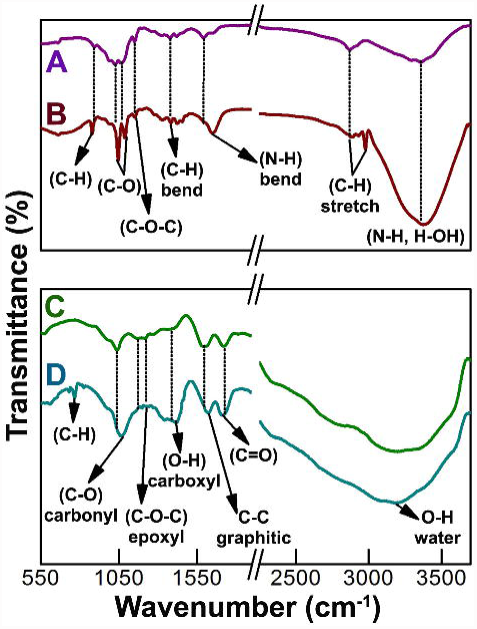
FTIR spectra of (A) pristine chitosan powder, (B) HNPs, (C) GO and (D) HNPs-GO.

The FTIR spectrum of GO samples (Fig. 5C) reveals the peaks relating to C-O (carbonyl) at 1040 cm^−1^, the C-O-C epoxyl group frequencies at 1175-1250 cm^−1^ and carboxyl-associated OH signal at 1405 cm^−1^ [14,32]. Well resolved peaks at 1600 and 1725 cm^−1^are assigned to the C-C vibrations of un-oxidized graphitic domains and C=O stretching vibrations, respectively [7,32]. The relatively broad peak centered at 3240 cm^−1^ is associated with the adsorbed water on the surface of the GO. As discussed previously, HNPs are expected to form chemical bonds at the basal planes and edges of GO. After functionalization with HNPs, the carbonyl peak of GO is broadened and shifted from 1040 cm^−1^ (Fig. 5C) to 1068 cm^−1^(Fig. 5D). The epoxyl group frequencies were almost dispersed and the peaks of carboxyl-associated OH, C-C graphitic domains and C=O stretching vibrations were also changed. Further, a new peak centered at 771 cm^−1^, along with two shoulder peaks (735 and 830 cm^−1^) attributed to C-H of chitosan, [15] was observed. This provides the supporting information for the functionalization of HNPs on the GO surface.

To gain further understanding of the chemical structure of pristine and hybrid nanostructures, ^1^H-NMR spectral analysis was utilized. Fig. 6A shows the ^1^H-NMR spectrum of HNPs, which exhibits the characteristic resonance peaks attributed to the functional groups of chitosan such as –C*H*–C*H*– (0.8 and 1.2 ppm), –N*H*– (1.9 ppm) and O*H* (5.3 ppm) [33]. Proton signals of GO nanostructures (Fig. 6B) are identified at 1.2, 4.5, 8.1 and 9.5 ppm and attributed to the –C*H*–C*H*–, O*H*, –C–COO*H*– and =C–COO*H*, respectively [33,34]. The spectrum of HNPs-GO (Fig. 6C) also shows the inherent –C*H*–C*H*– protons. The peak shift located at 1.9-2.0 ppm, attributed to the amine protons of chitosan, is relatively weaker than that of pristine HNPs (Fig. 6A), indicating the functionalization of HNPs on GO. Successful chemical bonding of HNPs on the oxygenated functional groups of GO are validated by the absence of free carboxyl proton signals (at 8.1 and 9.5 ppm) and appearance of multiple amide proton signals (at 6.5, 6.9, 7.1 and 7.2 ppm). The presence of reactive epoxyl and carboxyl groups on the GO lattice structures offered the necessary binding sites for the chemical decoration of HNPs.

**Fig. 6.**
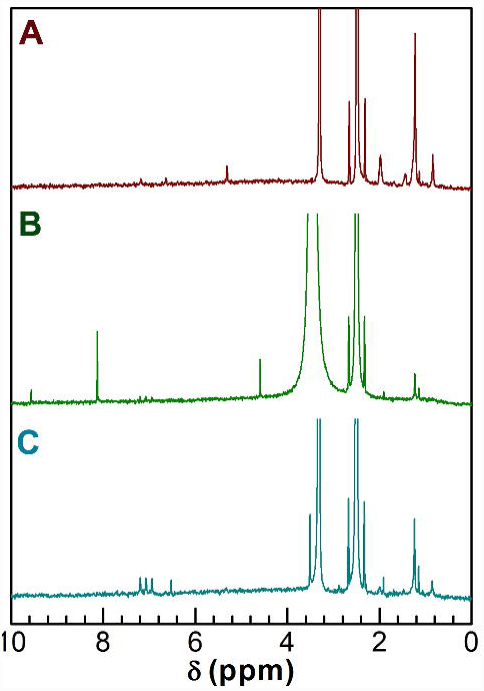
^1^H-NMR spectra of (A) HNPs, (B) GO and (C) HNPs-GO samples in DMSO-d_6_ solvent.

### 3.5 Raman spectroscopy

Raman spectral analysis further revealed the structural integrity of GO after chemical interaction with HNPs. The typical characteristics of Raman spectra of graphite materials are a G-band at 1570 cm^−1^ attributed to the *E*_2g_ phonon of sp^2^ C domains [35] and a D-band at 1345 cm^−1^ attributed to the vibrations of disordered C domains of graphite [32,35]. The presence of D-band at 1355 cm^−1^ and a G-band at 1596 cm^−1^ supports the oxygenation of graphite (Fig. 7). Chemically decorated HNPs on the surface of the GO lattice displayed a broadened D-band at 1355 cm^−1^ and G-band at 1583 cm^−1^ (red shifted from inherent 1596 cm^−1^), respectively. A slight change in the intensity ratio of the D- and G-bands (*I*_D_/*I*_G_) of HNPs-GO (0.85) compared to that in GO (0.81) indicated that functionalization of HNPs altered the in-plane sp^2^ graphitic domains of GO. According to an empirical formula known as the Tuinstra-Koenig relation, [36] the average crystallite size of the ordered graphitic sp^2^ C domains can be calculated using the following equation,

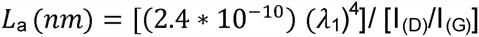

where *L*_a_ is the average crystallite size of the sp^2^ domains, λ_l_ is the input laser energy, *I*_(D)_ is the intensity of the D band, and *I*_(G)_ is the intensity of the G band. The calculated *L*_*a*_ values are 20.7 and 19.7 nm for GO and HNPs-GO, respectively. Observed changes in the size of sp^2^ hybridized domains are ascribed to the chemical interaction with HNPs. These results are in agreement with similar reports on GO hybridized with metal oxide nanoparticles [37] and biomaterials [38].

**Fig. 7.**
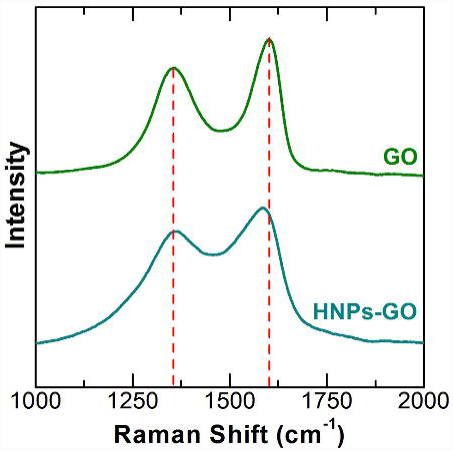
Raman spectra of GO and HNPs-GO.

### 3.6 Fabrication and electrochemical properties of HNPs-GO electrodes

Compared to conventional electrodes, carbon electrodes modified with conductive hierarchical nanostructures exhibit an enhanced electron transfer rate and more durable electrochemical properties [8,29]. Immobilization of bio-friendly conductive nanostructures with active chemical groups suitable for anchoring antibody or enzyme is certainly valuable for fabrication of label-free biosensor platforms [39]. To understand its feasibility as transducer material for an electrochemical biosensor platform, the primitive electrochemical response of the HNPs-GO was evaluated in comparison with pristine GO. Fig. 8 shows the pictorial representation of an integrated three-electrode system used for modification of pristine GO and HNPs-GO. Unlike conventional electrochemical systems, there is no external counter or reference electrodes utilized in the present study. In order to find an optimal potential region suitable for the prepared materials, a pre-screening CV measurement was performed between □1.0 to +1.0 V. From the analysis, it was found that □0.25 to +0.8 V is an optimal potential region for studying the redox behavior of the HNPs-GO modified Au-PCB electrodes.

**Fig. 8.**
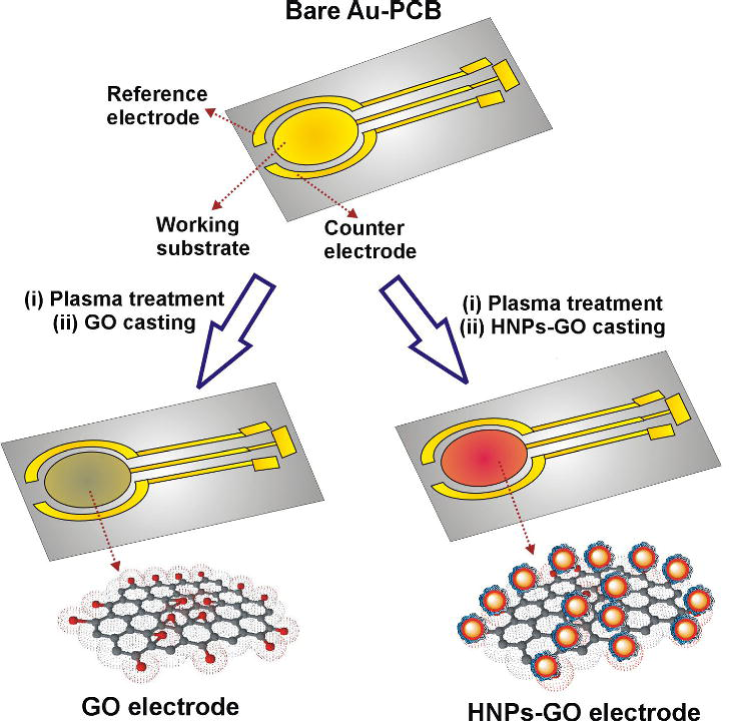
Schematic illustration of an integrated three-electrode system and modification with GO and HNPs-GO nanosheets.

Fig. 9A represents the CV curves of bare Au-PCB, pristine GO and HNPs-GO modified Au-PCB electrodes recorded at a constant scan rate of 50 mV/s. A 10 mM PBS solution containing the final concentration of 0.0027 M potassium chloride and 0.137 M sodium chloride, with a pH 7.4 was used as the supporting electrolyte. Under experimental conditions, bare Au-PCB and pristine GO electrodes don’t exhibit significant redox behavior. Pristine HNPs without GO as an interface layer are poorly stable on the Au-PCB substrate, which resulted in leaching and hindered the durable electrochemical response (data not shown). On the other hand, HNPs-GO modified electrode showed well-defined and highly amplified anodic peaks A_1_ at +0.38 V and A_2_ at +0.52 V; the former is related to the oxidation of Ag →Ag_2_O [40] and the latter is derived from the oxidation reaction of [Ru(bpy)_3_]^2+^→ [Ru(bpy)_3_]^3+^. Interestingly, the HNPs-GO electrode displayed a single cathodic peak at □0.12 V, suggesting a coherent reduction reaction of Ag_2_O→Ag and [Ru(bpy)_3_]^3+^ →[Ru(bpy)_3_]^2+^. The anodic peak current (A_2_: *I*_p_= +43 μA) generated from the oxidation reaction of [Ru(bpy)_3_]^2+^ is higher than the peak current of A_1_ (*I*_p_= +33 μA). This is probably attributed to the existence of [Ru(bpy)_3_]^2+^ on the surface of Ag core nanostructures which supported the overall metal-to-ligand charge transfer process. Further, it is speculated that the presence of chitosan layer on the core Ag@[Ru(bpy)_3_]^2+^ is expected to have reasonable influence on the observed redox wave.

**Fig. 9.**
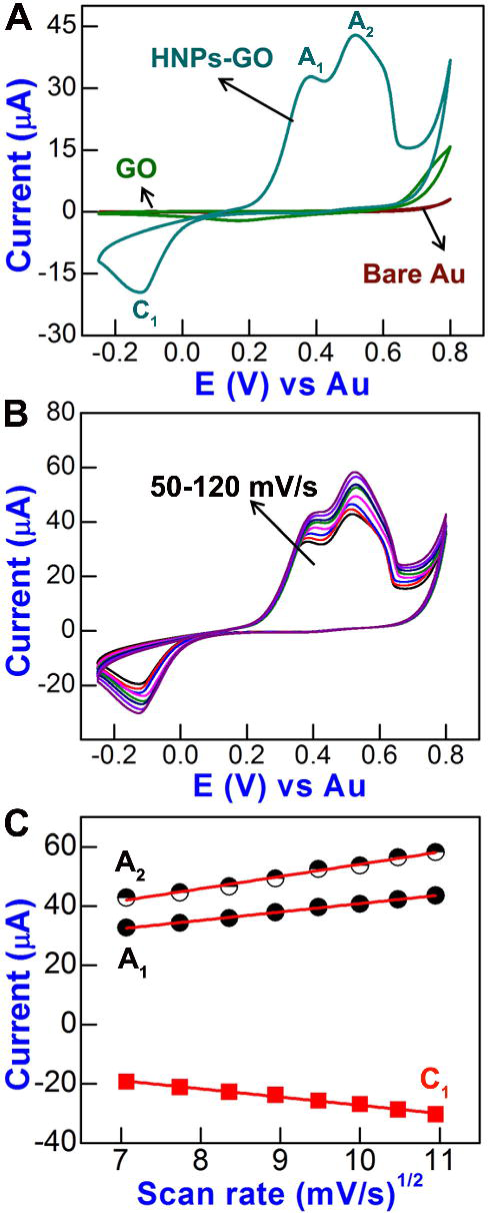
(A) CVs of bare Au-PCB, pristine GO and HNPs-GO modified electrodes. (B) CVs of HNPs-GO electrode at different scan rates (50-120 mV/s) in 10 mM PBS (pH 7.4) and (C) the corresponding plots of anodic (A_1_ and A_2_) and cathodic (C_1_) peak currents against the square root of scan rates.

Earlier studies demonstrated that the insulating nature of GO nanosheets could be transformed by functionalization of metallic composites, which not only provides better electrical conductivity but also creates a 3D hierarchical environment with a large surface area for rapid electron transfer. For instance, conductive polyaniline interconnected Fe_2_O_3_-rGO composites exhibited a surface-confined redox transition at the electrode interface [9]. Enhanced redox waves generated from Ag-doped with organometallic or conductive polymer composite electrodes have been reported previously in the literature [40,41]. Likewise, herein the hybrid combination of chemically interacted core Ag@[Ru(bpy)_3_]^2+^ and shell chitosan enabled a significant redox reaction at the GO interface. Such hybrid nanoplatform is useful for constructing advanced biosensor platforms.

In order to evaluate the constancy of the redox potentials and increasing peak currents in respect to the scan rate, CVs for HNPs-GO electrodes were recorded at different scan rates from 50 to 120 mV/s (Fig. 9B). The enhancement of the anodic (A_1_ and A_2_) and cathodic (C_1_) peak currents are in relation to the scan rate (Fig. 9C). The correlation coefficients for the anodic peaks were 0.9987 (*I*_p_A_1_), 0.9952 (*I*_p_A_1_) and the cathodic peak was 0.9964 (*I*_p_C_1_), indicating that it is a surface-confined process. This interesting redox behavior, which has emerged from the current investigation on CVs of HNPs-GO is valuable and provides the possibility of exploring their bio-affinity toward novel molecules, through a label-free, direct electrochemical detection strategy.

## 4. Conclusion

Three-dimensional nanosheets of GO decorated with HNPs composed of metal-dye complex (Ag@[Ru(bpy)_3_]^2+^) core and biopolymer (chitosan) shell was fabricated through a facile scalable wet-chemical approach. Chemically immobilized HNPs altered the inherent sp^2^-sp^3^ carbon domains on GO lattice and revealed functional changes in their optical and structural properties. Electron microscopic investigations supported the topographical information on distribution of HNPs on the ultra-thin sheets of GO. Electrodes modified with HNPs-GO showed an amplified and durable redox behavior compared to those of GO or HNPs. These findings suggest that hierarchical structures of HNPs-GO with multi-functional optical, electrochemical and biocompatible feasibilities are promising for further exploration in biosensing studies.

## Appendix A. Supplementary data

Supplementary data associated with this article can be found, in the online version, at

## Highlights

- Functionalization of hybrid Ag@[Ru(bpy)_3_]^2+^/chitosan on graphene oxide nanosheets
- Study of physico-chemical properties of graphene oxide-decorated with hybrid NPs
- Fabrication and electrochemical study of hybridized graphene oxide electrodes
- Feasibility of hybrid graphene oxide electrodes as an advanced biosensor platform

